## Supplemental Figures for "Enhanced specificity of high sensitivity somatic variant profiling in cell-free DNA via paired normal sequencing: design, validation, and clinical experience of the MSK-ACCESS liquid biopsy assay"

**Validation, and clinical experience of MSK-ACCESS, an ultra-deep cell-free DNA sequencing assay for non-invasive somatic genomic profiling**

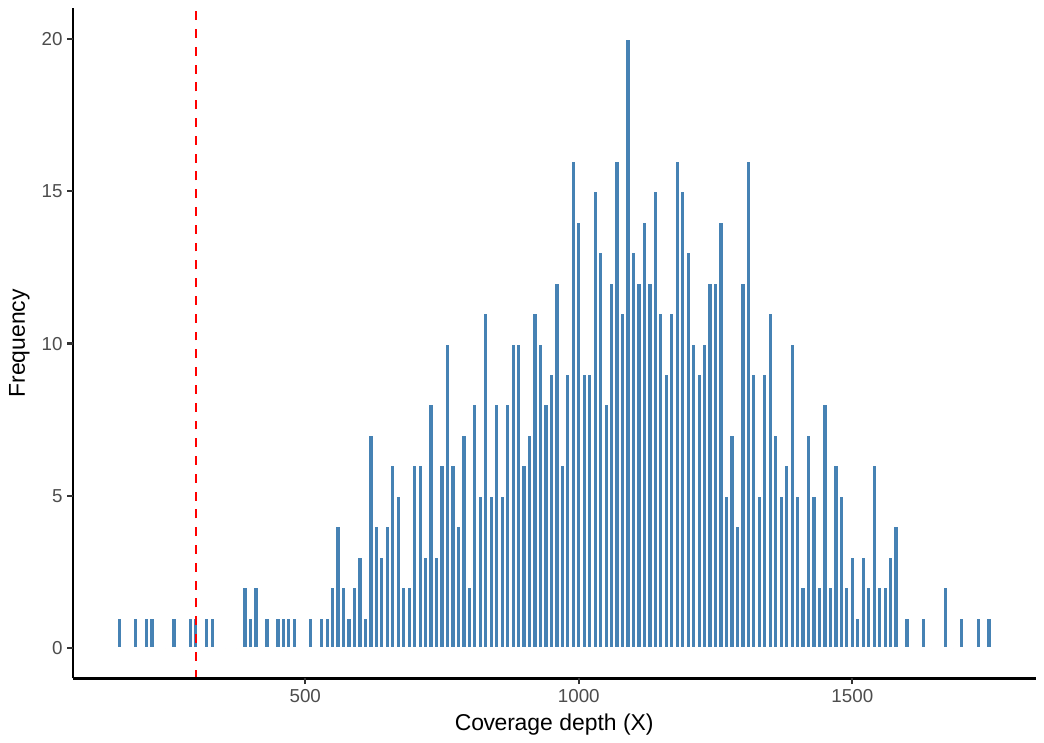

**Supplemental Figure 1: MSK-ACCESS exon coverage.** Mean coverage of all MSK-ACCESS exons across 47 healthy donor plasma samples.

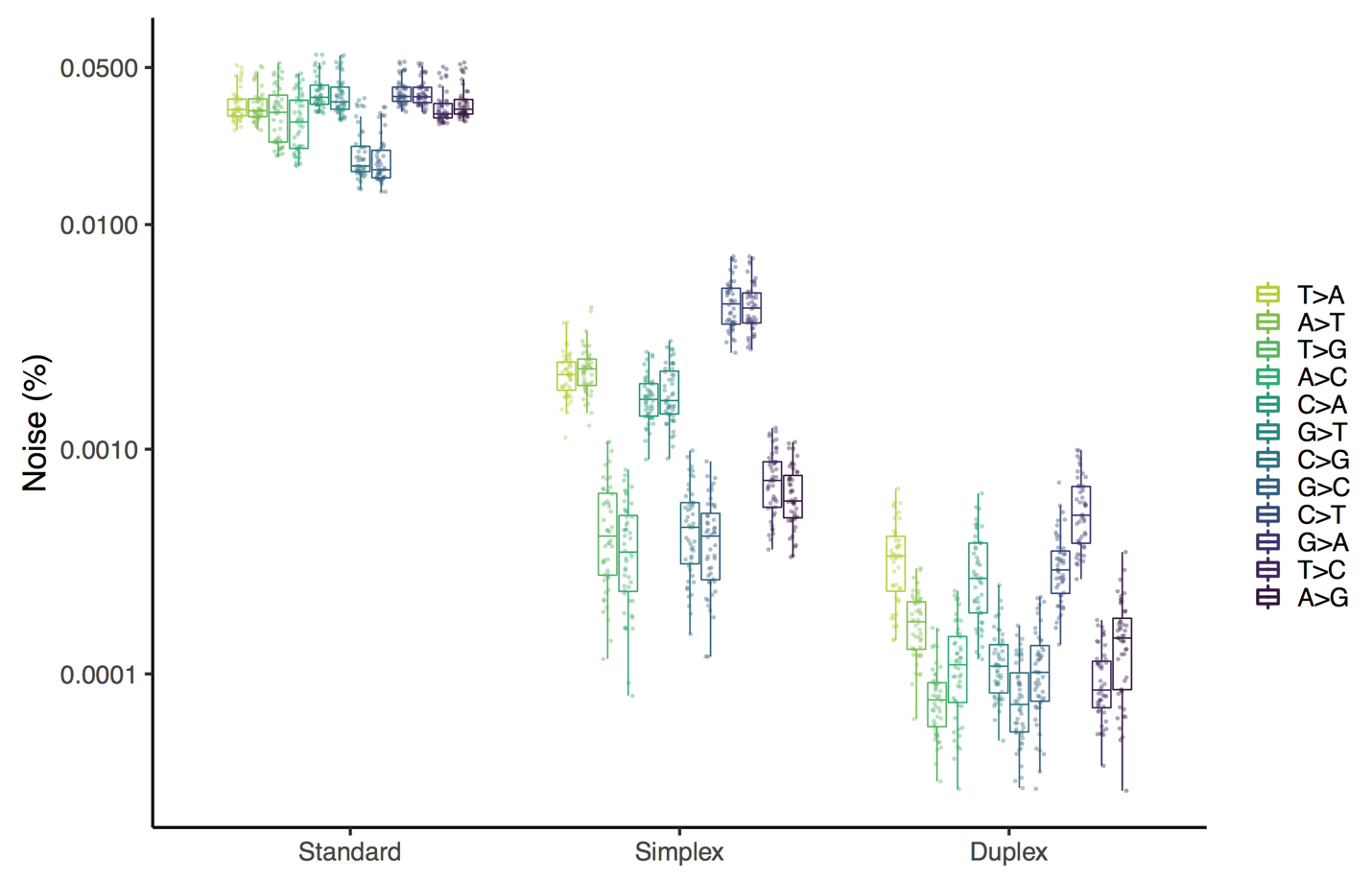

**Supplemental Figure 2: HiSeq 2500 error rate.** Error rate at sites with non-reference alleles in MSK-ACCESS panel in 47 healthy donor plasma samples sequenced on the HiSeq 2500.

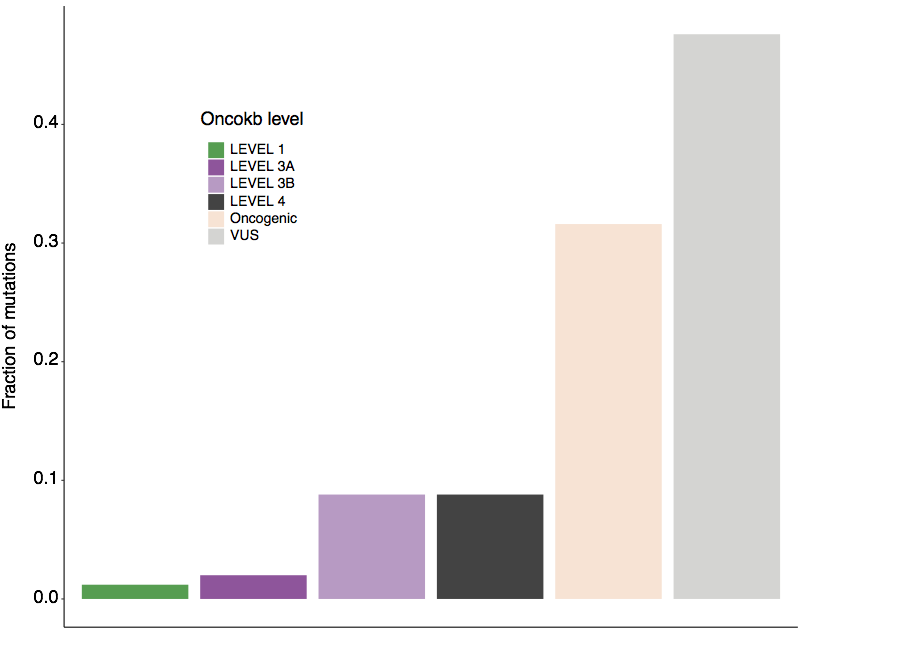

**Supplemental Figure 3: OncoKB levels of MSK-ACCESS-only mutations.** Fraction of mutations called only by MSK-ACCESS and colored by OncoKB designation

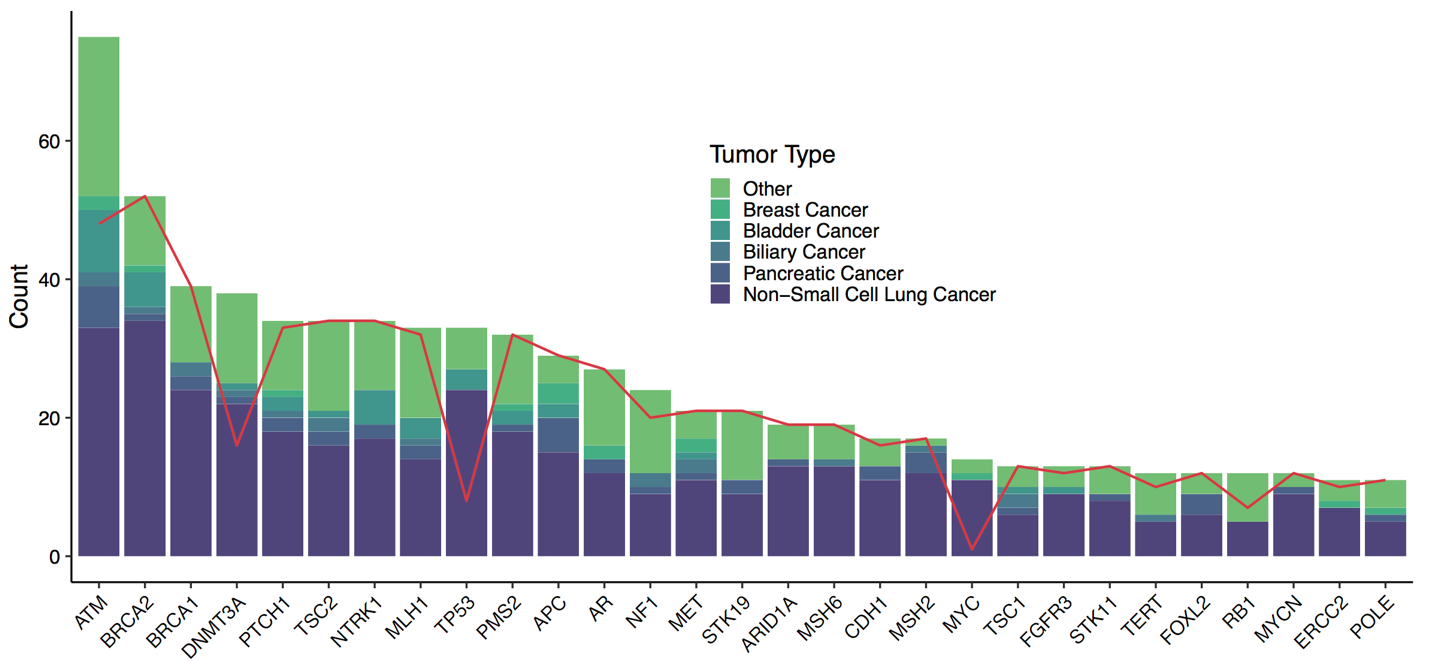

**Supplemental Figure 4: Genes filtered out with matched WBC.** Number of mutations per gene and tumor type that would remain in clinical samples after filtering with curated healthy normal plasma samples and gnomAD if WBC normal was not used. The red line denotes the number of total mutations that would be present if calls were further filtered for presence in COSMIC hematologic samples; while potential CH mutations would be removed, some tumor derived mutations would also be filtered out.

Patient plasma vs healthy normal plasma sample

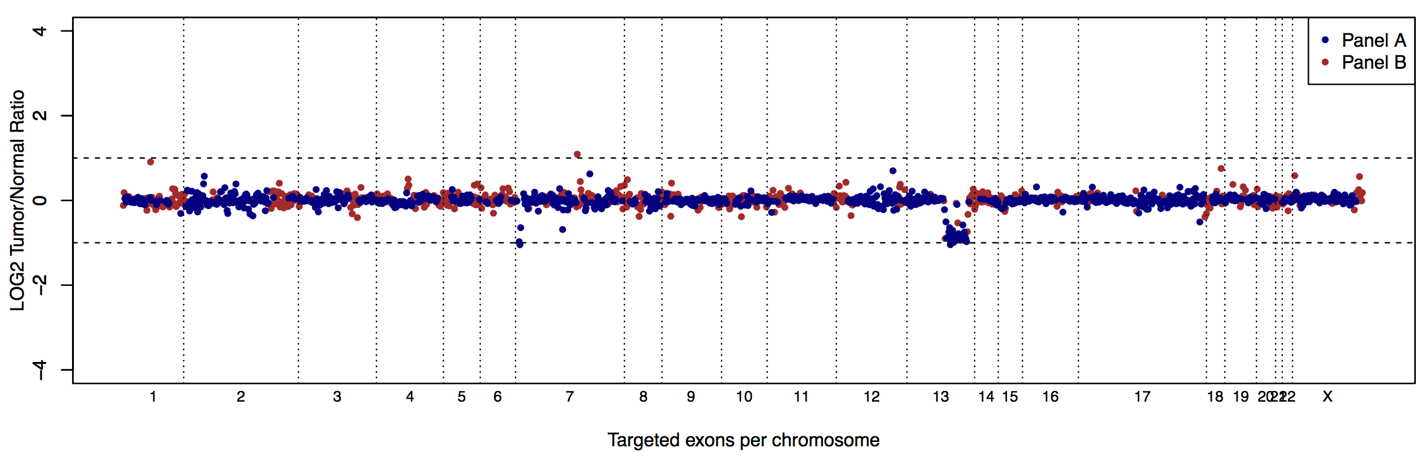

Patient WBC vs healthy normal WBC sample

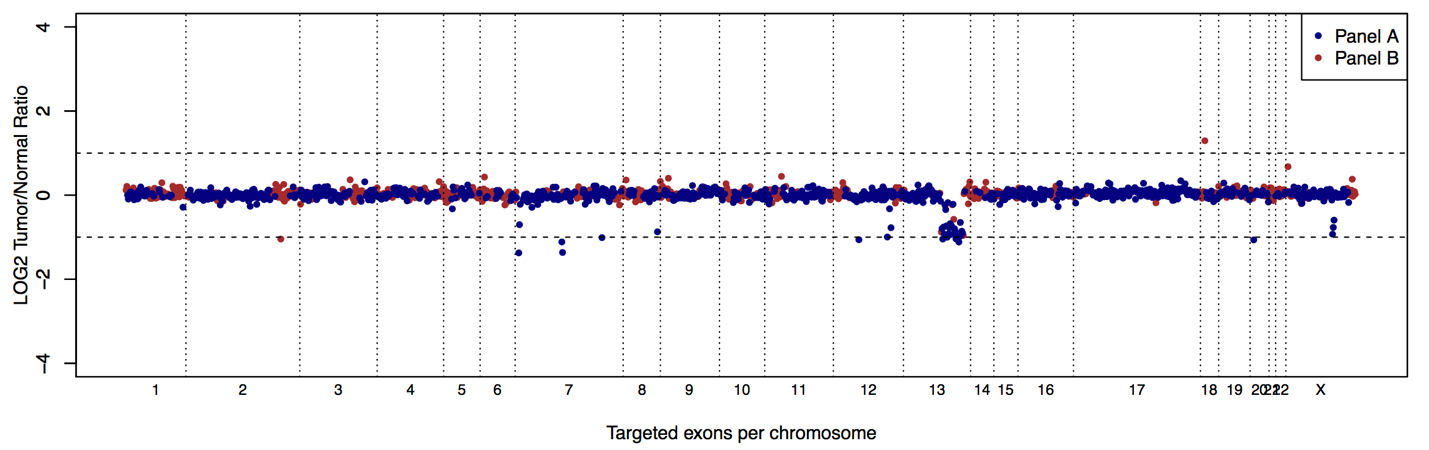

**Supplemental Figure 5: Germline copy number deletions in MSK-ACCESS.** In both the retinoblastoma cancer patient’s plasma and WBC, an *RB1* deletion was detected. Because WBC was sequenced, this deletion was not reported as somatic.

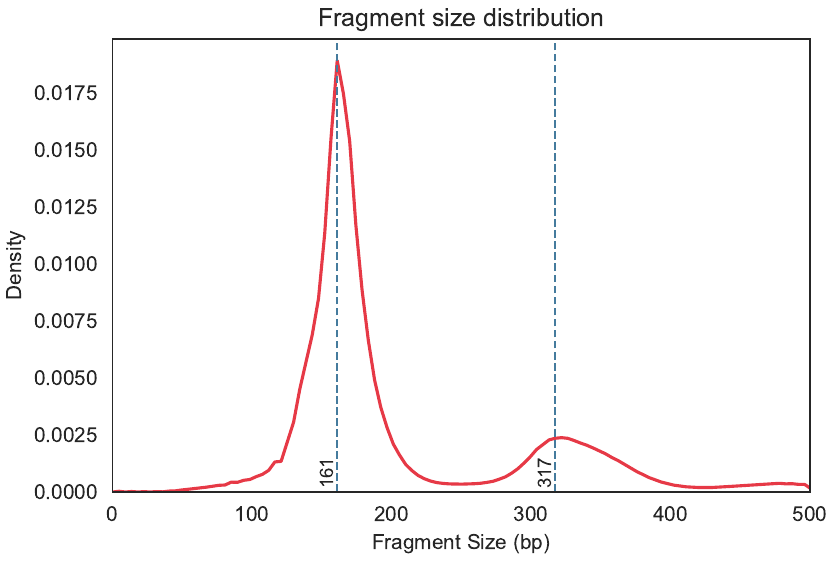

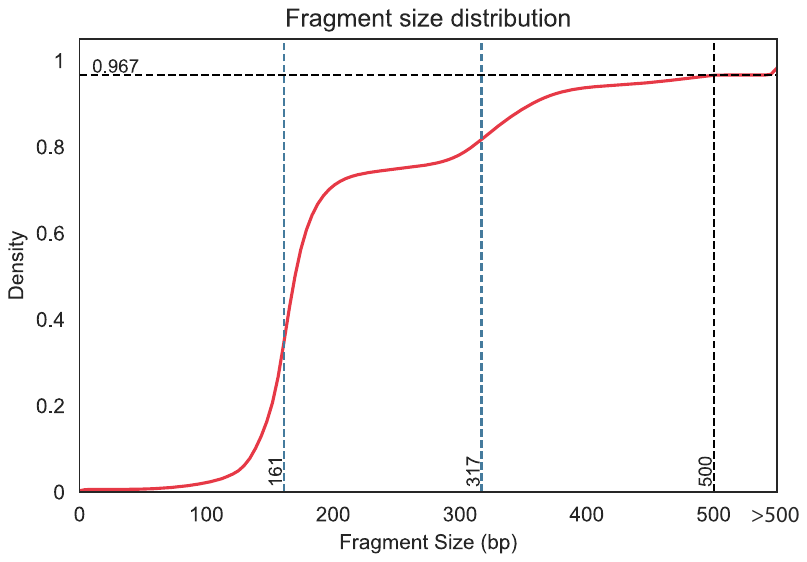

**Supplemental Figure 6. Fragment size distribution in plasma samples** (a) Probability density function plot indicating an expected bimodal distribution of fragment size with peaks at 161 bp and 317 bp. (b) Cumulative density function plot of the fragment size showing that fragments of sizes 500 bp or lower account for 96.7% of all fragments.
